# Integrin alpha 4 / beta 1 (CD49d/CD29) is a component of the murine IgG3 receptor

**DOI:** 10.1101/207274

**Authors:** Carolyn Saylor Hawk, Carolina Coelho, Diane Sthefany Lima de Oliveira, Verenice Paredes, Patrícia Albuquerque, Anamélia Lorenzetti Bocca, Ananésia Correa dos Santos, Victoria Rusakova, Heather Holemon, Maria Sueli Soares Felipe, Hideo Yagita, André Moraes Nicola, Arturo Casadevall

## Abstract

Antibodies exert several of their effector functions by binding to cell surface receptors. For murine IgG3 (mIgG3) the identity of its receptors (and the very existence of a receptor) is still under debate, as not all mIgG3 functions can be explained by interaction with Fcγ-receptor I (FcγRI). This implies the existence of an alternate receptor, whose identity we sought to pinpoint. We found that blockage of the alpha4/beta1 integrin (Itga4/Itgb1) selectively hampered binding of mIgG3 to macrophages and mIgG3-mediated phagocytosis. Manganese, an integrin activator, increased mIgG3 binding to macrophages. Blockage of FcγRI or Itgb1 inhibited binding of different mIgG3 antibodies to variable extents. Our results indicate an integrin component in the mIgG3 receptor. Given the more ancient origin of integrins in comparison with FcγR, this observation could have far ranging implications for our understanding of the evolution of antibody-mediated immunity, as well as in immunity to microorganisms, pathogenesis of autoimmune diseases and antibody engineering.

## Introduction

Antibodies are amongst the most studied biomolecules in history. Starting with late 19^th^ century studies on serum therapy that gave von Behring the first Nobel Prize in Medicine, a steady stream of important discoveries clarified how B-cells make antibodies, their structure and functions in both health and disease. This body of knowledge allowed harnessing of antibodies as biotechnological tools with major applications in diagnosis and therapy of human and animal diseases. Behind their success both as immune effectors and as biotechnological tools is the fact that antibodies are bifunctional molecules, with a variable domain that recognizes the antigen and a constant domain that mediates effector functions. For the IgG antibodies, the majority of these functions depend upon binding of the antibody Fcγ domain to cell-surface Fcγ-receptors (FcγR), which in turn activate (or inhibit) immune cells and regulate immunity (Casadevall and Pirofski, 2006; Nimmerjahn and Ravetch, 2008). Thus, binding of IgG to FcγR is essential to such varied phenomena as phagocytosis of a pathogen by macrophages, tissue damage resulting from deposited immune complexes and destruction of tumor cells by therapeutic anticancer antibodies.

The detection, cloning and sequencing of most FcγRs date from research in the 1970’s and 1980’s (Hulett and Hogarth, 1994). The latest discovered receptor, FcγRIV, was found by sequence homology with known FcγR α subunits (Nimmerjahn et al., 2005). The known FcγRs explain functions of murine IgG1, IgG2a and IgG2b. In spite of several decades of research, one major riddle remains in this field. In 1981, Diamond and Yelton posited that murine IgG3 (mIgG3) had its own receptor based on the observation that a clone of the macrophage-like J774 cell line lost the ability to phagocytose mIgG3-opsonized particles while retaining functionality with all other mIgG isotypes, indicating the existence of a distinct receptor for mIgG3 (Diamond and Yelton, 1981). In the late 1990s, another group reported that murine FcγRI, the high-affinity FcγR, was responsible for mIgG3 function, based on the failure of FcγRI-deficient murine bone marrow-derived macrophages (BMMs) to internalize mIgG3-coated erythrocytes (Gavin et al., 1998). However, subsequent studies have established that mIgG3 does not have measurable affinity for the classical FcγRs (Bruhns et al., 2009; Nimmerjahn and Ravetch, 2006, Nimmerjahn and Ravetch 2008). Furthermore, mIgG3 was found to be opsonic even when all the known FcγRs were absent or blocked (Saylor et al., 2010). In other models, mIgG3 antibodies mediated its function solely by activating complement, without evidence of binding to any cell-surface receptor (Azeredo da Silveira et al., 2002; Han et al., 2001; Hjelm et al., 2005). Thus, the receptor whose function was missing in Diamond’s 1981 sub-clone of the J774 cell line remains obscure, and the very existence of a mIgG3 receptor is uncertain.

In human cells, all human IgG isotypes unambiguously interact with known FcγRs (Hulett and Hogarth, 1994; Nimmerjahn and Ravetch, 2006, 2008). There is no direct equivalence between murine and human IgG isotypes, but human IgG2 (hIgG2) is related to mIgG3 because both mice and humans have a clear bias towards responding to carbohydrate antigens with these isotypes (Perlmutter et al., 1978; Scott et al., 1988; Slack et al., 1980). Additionally, mIgG3 frequently behaves as a cryoglobulin, being involved in autoimmune diseases such as glomerulonephritis or lupus-like skin lesions in mice (Abdelmoula et al., 1989; Kuroda et al., 2005; Otani et al., 2012); hIgG2 association with cryoglobulinemia in humans was never detected, though. Technological interest in the mIgG3 isotype is marginal because mIgG3 monoclonal antibodies are prone to aggregation as a result of inter-molecular constant region interactions (Greenspan and Cooper, 1992), which decreases productivity, stability and safety. However, engineering antibodies to modify their binding to different FcγRs and thus modulate pharmacological effects is an important strategy in monoclonal antibody drug development (Saxena and Wu, 2016). The existence and identity of this yet unknown non-FcγR IgG receptor could thus be relevant in Immunology, Immunopathology and production of therapeutic antibodies.

Here we show the results of a loss-of-function screening with shRNAs to 15,000 murine genes indicating that integrin beta 1 (Itgb1) and integrin alpha 4 (Itga4) form a cell-surface receptor for mIgG3 in mouse macrophages. We provide experimental validation of this finding via loss and gain of function assays for several mIgG3 antibodies. The finding of a non-FcγR receptor for IgG antibodies could have large impact on our understanding of the humoral immune response and on antibody engineering.

## Results

### Screening for the IgG3 receptor using pooled shRNA libraries

We designed and conducted a screen for the mIgG3 receptor in mice with a pooled shRNA library from the RNAi Consortium (Moffat et al., 2006). Using engineered lentiviral vectors targeting 15,000 different mouse genes we transduced J774 cells to generate a pool of macrophage-like cells, each with a single gene knocked down. We screened the knock-down library by sorting twice for cells that showed decreased binding to mIgG3 but had normal levels of mIgG1 binding (Figure 1A-B). Sequencing the shRNA sequences of pre-sort and post-sort J774 pools determined that 8,145 of the 8,937 shRNA sequences detected were differentially represented in the separated populations. Of these, 2,794 had higher representation in the post-sort population, i.e. low binding to mIgG3. We narrowed our search to select for plasma membrane proteins, which were targeted by 266 of these shRNAs. Among these 266, we chose five candidate genes, based on known macrophage expression and possible role in phagocytosis or Ab binding: Lrig2 (leucine-rich repeats and immunoglobulin-like domains 2), Itgb21 (Integrin beta 2-like, or Pactolus), Itgb1 (Integrin beta 1), Tlr2 (toll-like receptor 2), and Fasl (Fas ligand).

**Figure 1.**
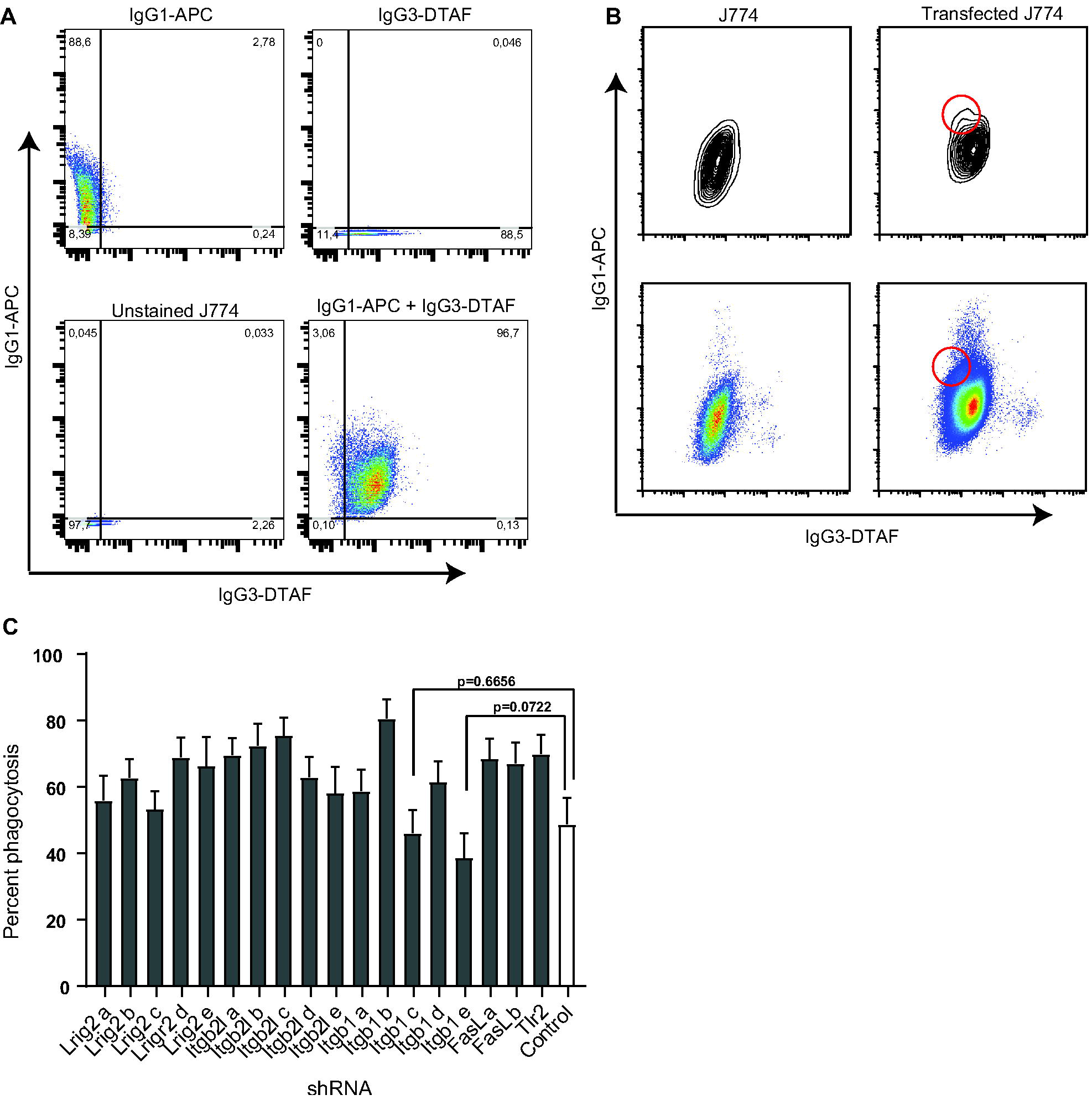
High-throughput screening for a mIgG3 receptor. Panels A and B show J774 cells transduced with the entire library and stained with fluorescent mIgG1 and mIgG3, whereas panel C shows a secondary screen with individual shRNA clones to test some candidates. (A) Untransduced J774 cells stained with mIgG1-APC and/or mIgG3-DTAF. (B) Comparison of untransduced J774 cells and J774 cells transduced with shRNA library, stained with both mIgG1-APC and mIgG3-DTAF. Data from bottom two panels are the same as the top two panels represented differently to better visualize the population of interest (approximated by red circle). (C) Phagocytosis assay using *C. neoformans* opsonized with 3E5-mIgG3 and J774 cells transduced with 17 individual shRNA-encoding lentiviruses targeting five candidates from the screening. All cells were pre-incubated with CR block (mAbs specific for CD18, CDllb and CDllc) and Fc block (mAb 2.4G2, specific for FcγRII and FcγRIII) prior to addition of *C. neoformans*. Control cells were transduced with a non-target shRNA. Four fields were analyzed per well, with one well for each condition. Bars indicate percent phagocytosis (number of macrophages containing ingested *C. neoformans* divided by the total number of macrophages) with 95% confidence interval.

To confirm their involvement in mIgG3 binding, we transduced individual shRNA sequences targeting these candidates (5 each for Lrig2, Itgb2l and Itgb1, 2 for Fasl and 1 for Tlr2) into J774 cells. We then performed phagocytosis assays to detect decreased uptake of *C. neoformans* after opsonization with 3E5-IgG3, a hybridoma-derived monoclonal antibody that binds to the fungal capsule (Yuan et al., 1995). The experiment was performed in the presence of 2.4G2 antibody (Fc block) - which blocks CD16 and CD32, the alpha subunits of FcγRIII and FcγRII. We also used antibodies to CD18, CD11b and CD11c (CR block) to inhibit complement receptors CR3 and CR4, which can directly bind to *C. neoformans* capsule carbohydrate upon antibody binding to the capsule and promote complement independent phagocytosis (Taborda and Casadevall, 2002). Although all candidates showed uptake of mIgG3-*C. neoformans*, two clones transduced with *Itgb1* shRNAs showed lower (albeit not statistically significant) phagocytic efficacy (Figure 1C). These two clones, Itgb1 c. and e., were also the ones that showed the greatest reduction in surface expression of Itgb1 (Figure SI).

### mIgG3 binding to macrophages depends on Itgb1 expression and divalent cations

Given the encouraging screening hit, we tested the effect of integrins on binding of fluorescently labeled mIgG3 and mIgG1 antibodies to macrophages. First we took advantage of the fact that integrin activity is modulated by divalent cations (Leitinger et al., 2000) to test antibody binding to macrophages in the presence or absence of Mg^2+^, Mn^2+^ and Ca^2+^. Consistent with an integrin type of interaction, 1 mM Mn^2+^ increased the binding of mIgG3 to J774 cells. In the presence of 1 mM Ca^2+^ and of 1 mM Mg^2+^ mIgG3 binding was increased and unchanged, respectively (Figure 2A). We next tested a range of concentrations of each cation, which showed that the increased mIgG3 binding resulting from supplementation with divalent cations was dose-dependent and that the divalent cations had little effect on mIgG1 binding to cells (Figure S2). Finally, when these cations were combined, the increased mIgG3 binding in the presence of Mn^2+^ and Ca^2+^ was additive, whereas addition of Mg^2+^ had little effect on Ca^2+^ but dampened the effect of Mn^2+^ (Figure S3).

**Figure 2.**
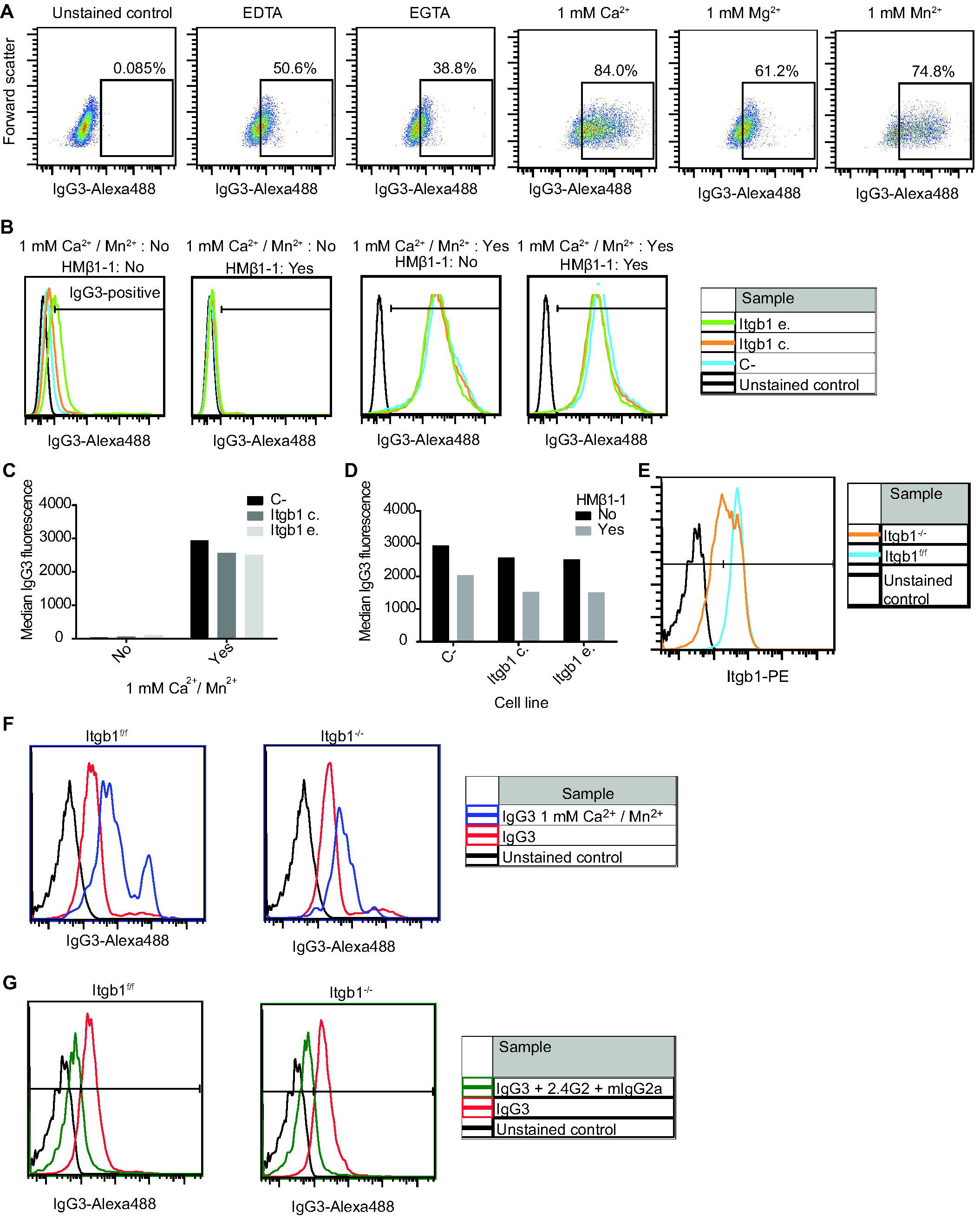
Divalent cations and *Itgb1* expression affect mIgG3 binding to macrophages. (A) Fluorescently labeled 3E5-mIgG3 was used to measure by flow cytometry binding of mIgG3 to J774 cells in presence of the divalent cation chelator EDTA, the Ca^2+^-specific chelator EGTA and the cations Ca^2+^, Mg^2+^ and Mn^2+^. All chelators and salts were present at 1 mM in PBS. (B) Fluorescently labeled 3E5-mIgG3 was used to measure by flow cytometry binding of mIgG3 to J774 cells transduced with Itgb1 and control shRNA in presence or absence of HMβ1-1. (C and D) Median intensity fluorescence of some of the samples depicted in panel B, showing that in the presence of ImM Ca^2+^ / Mn^2+^ Itgb1 shRNA-mediated knockdown decreases IgG3 binding. (D) Median intensity fluorescence of other samples depicted in panel B showing that both in the presence or absence of ImM Ca^2+^/ Mn^2t^, HMβ1-1 decreases mIgG3 binding. (E) Itgb1 expression on *Itgb1*^−/−^ or *Itgb1*^f/f^ BMM, detected by flow cytometry with fluorescently labeled HMβ1-1. (F) Binding of fluorescently labeled 3E5-mIgG3 to BMM in the presence of ImM Ca^2+^ / Mn^2+^. (G) Binding of fluorescently labeled mIgG3 to BMM in presence of excess monomeric mIgG2a.

Next we evaluated the binding of IgG3 to J774 cells under *Itgb1* interference by shRNA and/or HMβ1-1, a hamster mAb that inhibits murine Itgb1. To increase the efficiency of *Itgb1* knockdown, we sorted the population of J774 cells transduced with Itgb1 c. and e. and selected for cells with lowest expression of Itgb1. With integrins activated by Mn^2+^ and Ca^2+^, *Itgb1* knockdown slightly decreased mIgG3 binding in comparison with control cells (Figure 2B-C). In the presence of HMβ1-1 all three cell lines manifested decreased binding of IgG3 (Figure 2D). When the effects of the shRNAs and HMβ1-1 were combined in the presence of divalent cations, the median IgG3 fluorescence decreased by roughly 50%.

To confirm the results obtained with macrophage-like J774 cells, we used primary macrophages. Since *Itgb1* is essential in many cellular functions, genetically deficient (knock-out) mice are not viable. To overcome this, we used BMM from LysM-Cre, Itgb1^flox/flox^ (Itgb1^−/−^) mice, in which *Itgb1* is conditionally knocked out in macrophages and other myeloid cells. Flow cytometry with PE-labeled HMβ1-1 showed *Itgb1* deletion in about 50% of the BMM Itgb1^−/−^ mice, in comparison with a 95% Itgb1-positive population derived from the Cre-negative, Itgb1^flox/flox^ (Itgb1^f/f^) control littermate mice (Figure 2E). Flow cytometric mIgG3 binding assays showed that like J774 cells, BMM bound monomeric mIgG3 and that binding was increased in presence of Ca^2+^ and Mn^2+^. However, binding of Ab to the cells was similar between Itgb1^−/−^ and Itgb1^f/f^ cells (Figure 2F). Since BMM had been used by Gavin and colleagues (Gavin et al., 1998) to conclude that FcγRI is an mIgG3 receptor, we repeated the mIgG3 binding in presence of excess monomeric mIgG2a, which binds to FcγRI with high affinity (Nimmerjahn and Ravetch, 2008). This resulted in no clear difference between the Itgb1^−/−^ and Itgb1^f/f^ cells in mIgG3 binding to BMM (Figure 2G).

### Loss of Itgb1 function decreases mIgG3-mediated phagocytosis

Next we studied the role of Itgb1 in mIgG3-mediated phagocytosis. We repeated the phagocytosis assay with the sorted Itgb1 c. and Itgb1 e. knock-downs as well as cells transduced with a control shRNA; we also further interfered with Itgb1 function using HMβ1-1. As shown in Figure 3A, shRNA-mediated knockdown and HMβl-l blockage affected phagocytosis mediated by 3E5-mIgG3; in addition to decreasing the percent phagocytosis, *Itgb1* shRNA and HMβ1-1 resulted in a significant decrease in the number of *C. neoformans* cells internalized per macrophage (Figure 3B). Remarkably, we observed almost complete abrogation of *C. neoformans* phagocytosis mediated by 3E5-mIgG3 when both Itgb1 blocking strategies were combined.

**Figure 3.**
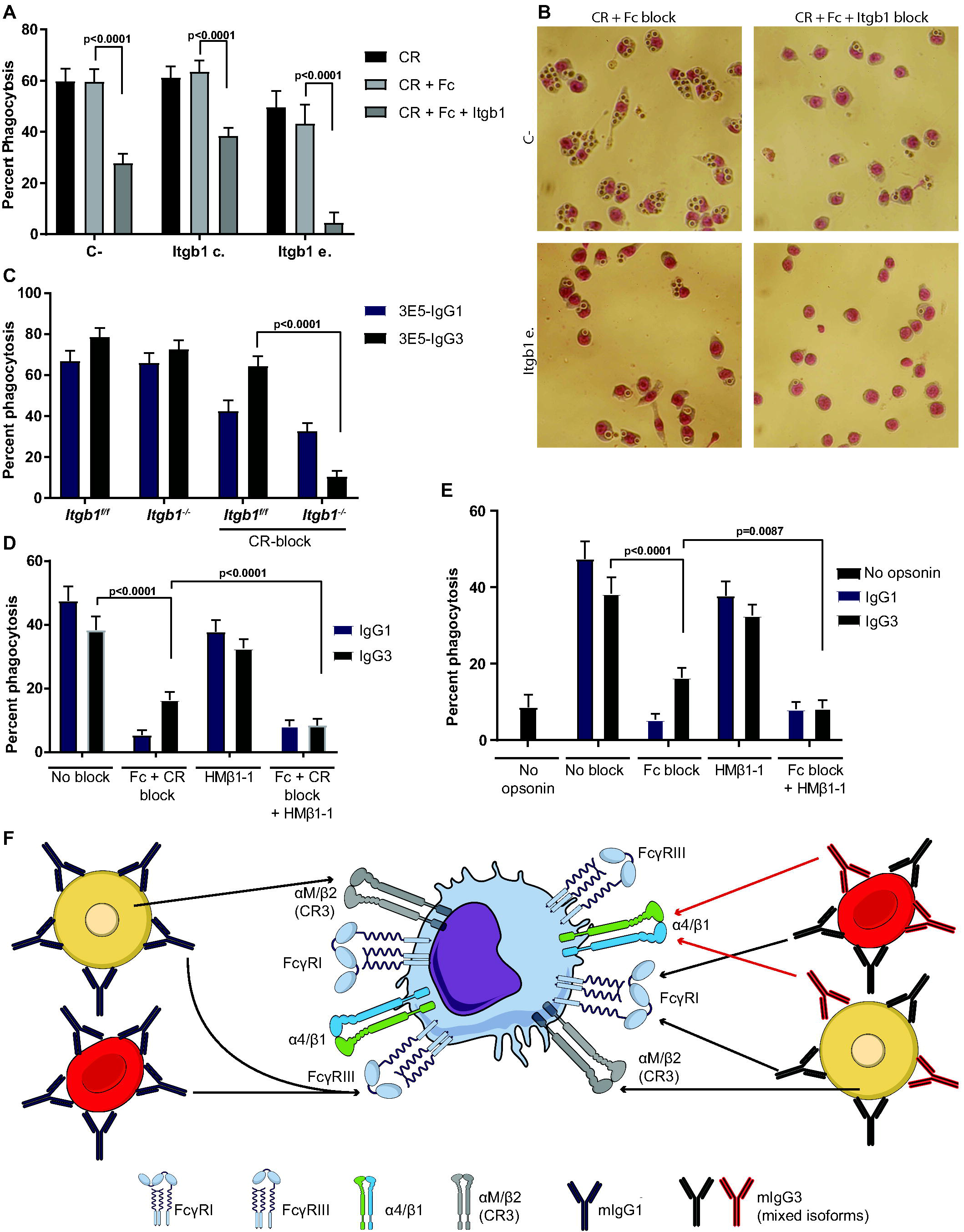
*Itgb1* silencing, knockout or blockage with HMβ1-1 decrease phagocytosis in macrophages. (A) Phagocytosis assay using *C. neoformans* cells opsonized with 3E5-IgG3 and Itgb1 c. and e. transduced J774 cell lines. Macrophages were pre-incubated with antibodies to block FcγRII/III and complement receptors, as indicated, prior to addition of fungi. (B) Images showing representative fields of the phagocytosis assay shown in panel A. Top two panels represent control cells (C- shRNA) and bottom two panels represent the Itgb1 e. transduced cell line, with blocking conditions denoted at top. (C) Phagocytosis assay with BMMs derived from *Itgb1*^−/−^ mice or their littermate Itgb1^f/f^ controls in the presence or absence of CR blocking Abs and *C. neoformans* cells opsonized with 3E5-mIgG3 or 3E5-mIgG1. Bars indicate the percent phagocytosis and the 95% confidence interval. (D) Phagocytosis assay with recombinant 2H1-mIgG1 and 2H1-mIgG3 antibodies and *C. neoformans*. Bars indicate the percent phagocytosis and the 95% confidence interval. (E) Phagocytosis of FITC-labeled sRBCs coated with recombinant 4-4-20-mIgG1 and 4-4-20-mIgG3 antibodies. In panels A, C, D and E the p-values were calculated with Fisher’s exact test. Bars indicate the percent phagocytosis and the 95% confidence interval. At least three fields were analyzed per well, with at least two wells per condition. (F) Model summarizing the data obtained with different phagocytosis assays. The black arrows indicate results that reproduced what had been previously known about the interaction of mIgG isotypes and Fcγ-receptors (Bruhns, 2012; Nimmerjahn et al., 2015) and between the *C. neoformans* capsule and CR3 (Taborda and Casadevall, 2002). The red arrows indicate the Itga4/Itgb1 role in mIgG3-mediated phagocytosis demonstrated in the other panels.

To confirm the results obtained with macrophage-like J774 cells, we used primary macrophages. BMM from both Itgb1^f/f^ and the *Itgb1*^−/−^ mice showed high levels of phagocytosis of *C. neoformans* cells coated with both 3E5-mIgG1 and 3E5-mIgG3. We had previously shown that BMM phagocytosis of 3E5-mIgG3-coated *C. neoformans* was not affected by Fc block and was only partially reduced by CR block, an indication of the existence of a non-Fcγ receptor (Saylor et al., 2010). In the present experiments, BMM from *Itgb1*^−/−^ mice phagocytosed *C. neoformans* to the same extent as those from control *Itgb1*^f/f^ mice, presumably because of the redundancy between Itgb1 and complement receptors (Figure 3C). In presence of CR block, however, 3E5-mIgG3-mediated phagocytosis was drastically reduced in *Itgb1*^−/−^ cells in comparison with control *Itgb1*^f/f^ BMM; integrin knockout had very little effect on 3E5-mIgG1 opsonization.

To test how generalizable these findings are, we used genetic engineering to produce two pairs of recombinant mIgG1 and mIgG3 antibodies. For the first pair, 2H1-mIgG1 and 2H1-mIgG3, we have used the variable region sequences of another antibody to *C. neoformans* capsule whose Fab structure has been solved by X-ray crystallography (Young et al., 1997). The other pair, 4-4-20-mIgG1 and 4-4-20-mIgG3, has the variable domains of a high affinity antibody to the hapten fluorescein isothiocyanate (FITC). Detailed structural information is also available for the 4-4-20 Fab structure (Whitlow et al., 1995). Recombinant antibodies were produced by co-transfecting mammalian cells with vectors expressing 2H1 and 4-4-20 heavy and light chains. Comparing the phagocytosis assays performed with *C. neoformans* cells coated with 3E5 hybridoma antibodies with the ones performed with 2H1 antibodies, we have observed the same pattern of almost complete abrogation of mIgG1-mediated phagocytosis upon CR and Fc block (Figure 3D). However, 2.4G2 did significantly reduce the phagocytosis of *C. neoformans* coated with 2H1-mIgG3 from 38% to 16% (p< 0.0001) when compared to mIgG3-coated *C. neoformans* in presence of CR block. This already low figure was further reduced to 8.4% upon addition of HMβ1-1 (p< 0.0001).

With the 4-4-20 antibody pair, we made phagocytosis assay using a different particle, FITC-labeled sheep red blood cells (sRBC). The background in this experiment was higher than that in experiments with *C. neoformans*, as we observed 8.7% phagocytosis with non-opsonized FITC-labeled sRBCs (Figure 3E). Addition of 2.4G2 reduced 4-4-20-IgG1 phagocytosis to the same level observed with non-opsonized sRBC. It also reduced 4-4-20-IgG3 phagocytosis, but to a level (19%) that was still significantly above background (p < 0.0001). HMβ1-1 alone had no effect on the phagocytosis rates. When 2.4G2 and HMβ1-1 were combined, the phagocytosis rate was decreased to 8.4% (p = 0.0087). The results from all phagocytosis experiments are summarized in Figure 3F.

### Integrin alpha 4 pairs with Itgb1 to mediate mIgG3 function

Integrins are heterodimeric complexes, composed of beta and alpha subunits. Using shRNAs to decrease surface expression of five alpha integrins that can pair with *Itgb1* and are known to be expressed in macrophages (*Itga3*, *Itga4*, *Itga5*, *Itga9* and *Itgav*), we tested which ones decreased phagocytosis of mIgG3-coated *C. neoformans*. As shown in Figure 4, two shRNAs targeting *Itga4* reduced 3E5-mIgG3 phagocytosis about two-fold, whereas the knockdown of any of the other alpha subunits had no effect on fungal internalization. We note that both Itgb1 and Itga4 knockdown had significant effects in cell health, an unavoidable confounding factor. However, we did detect loss of 3E5-mIgG3-mediated phagocytosis without loss of mIgG1 function for one of the shRNA clones targeting Itga4. Therefore, mIgG3 phagocytosis is dependent on Itgb1 and Itga4 surface expression, presumably due to integrin heterodimer formation.

**Figure 4.**
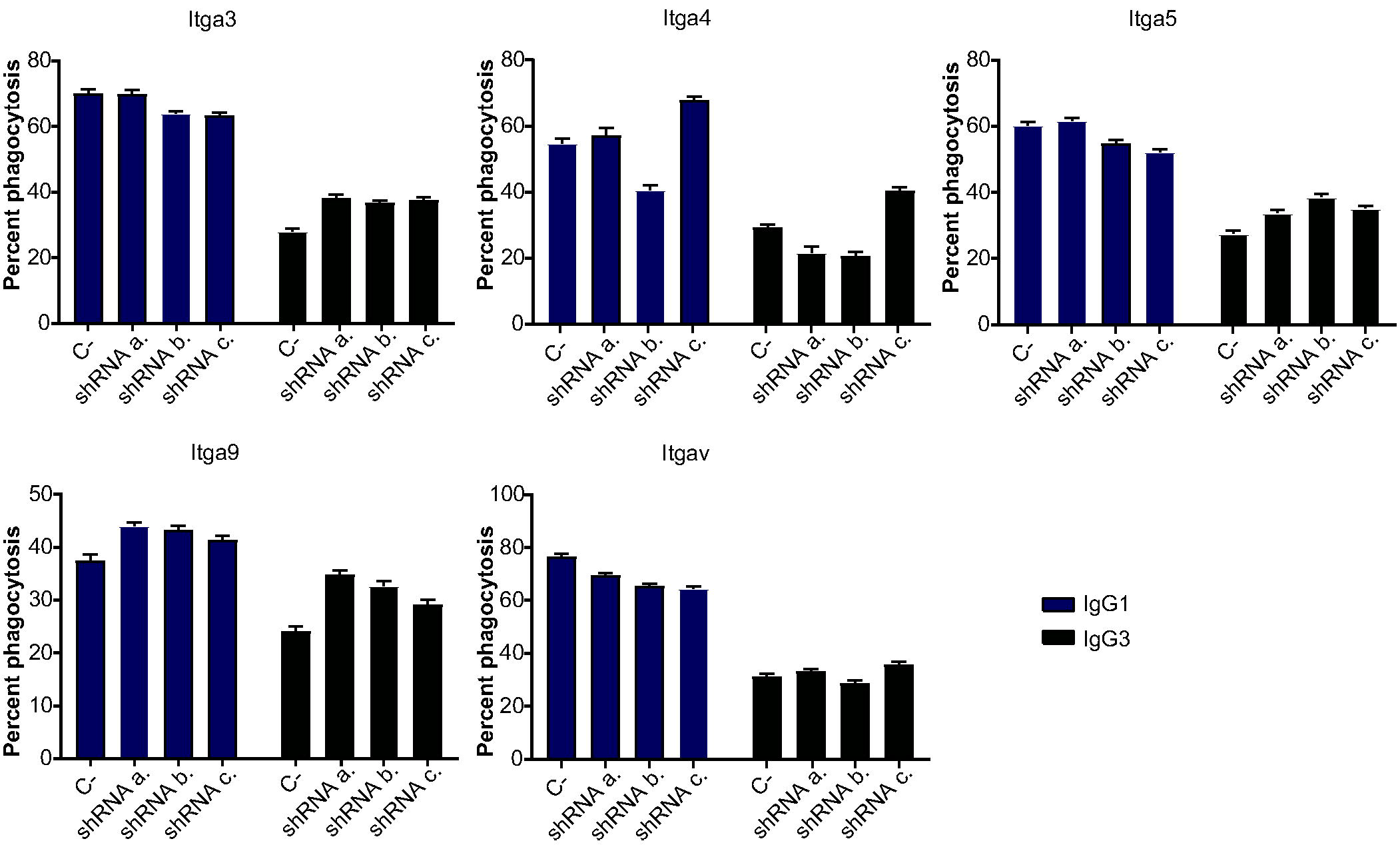
Phagocytosis with J774 cells transduced with shRNAs to integrin alpha chains. J774 cells were transduced with shRNAs targeting five integrin alpha chains which pair with Itgb1 and stably transfected cells selected with puromycin. A phagocytosis assay was then performed with 3E5-IgG1 or 3E5-IgG3 antibodies, in the presence of CR block, and fluorescently labeled *C. neoformans* using a scanning cytometer. Bars represent the percent phagocytosis and 95% confidence interval from one to two independent experiments, each containing three individual wells from which ^~^2,000 - 18,000 macrophages were evaluated. All p-values were calculated with Fisher’s exact test.

### Gain-of-function assays with *Itgb1*

We transfected Chinese Hamster Ovary (CHO) cells so they would express murine *Itgb1* to perform gain-of-function assays. In absence of divalent cations, binding of 3E5-mIgG3 but not 3E5-mIgG1 increased when comparing *CHO-Itgb1* cells with untransfected controls (Figure 5A). Similar to J774 cells, Ca^2+^ and Mn^2+^ increased 3E5-mIgG3 binding to CHO cells and murine *Itgb1* overexpression increased the proportion of 3E5-mIgG3-positive CHO cells from 75% to 97%. This increase in binding was almost completely abrogated by addition of HMβl-l, which does not bind to or inhibit native hamster integrin beta 1 on CHO cells (Figure 5B).

**Figure 5.**
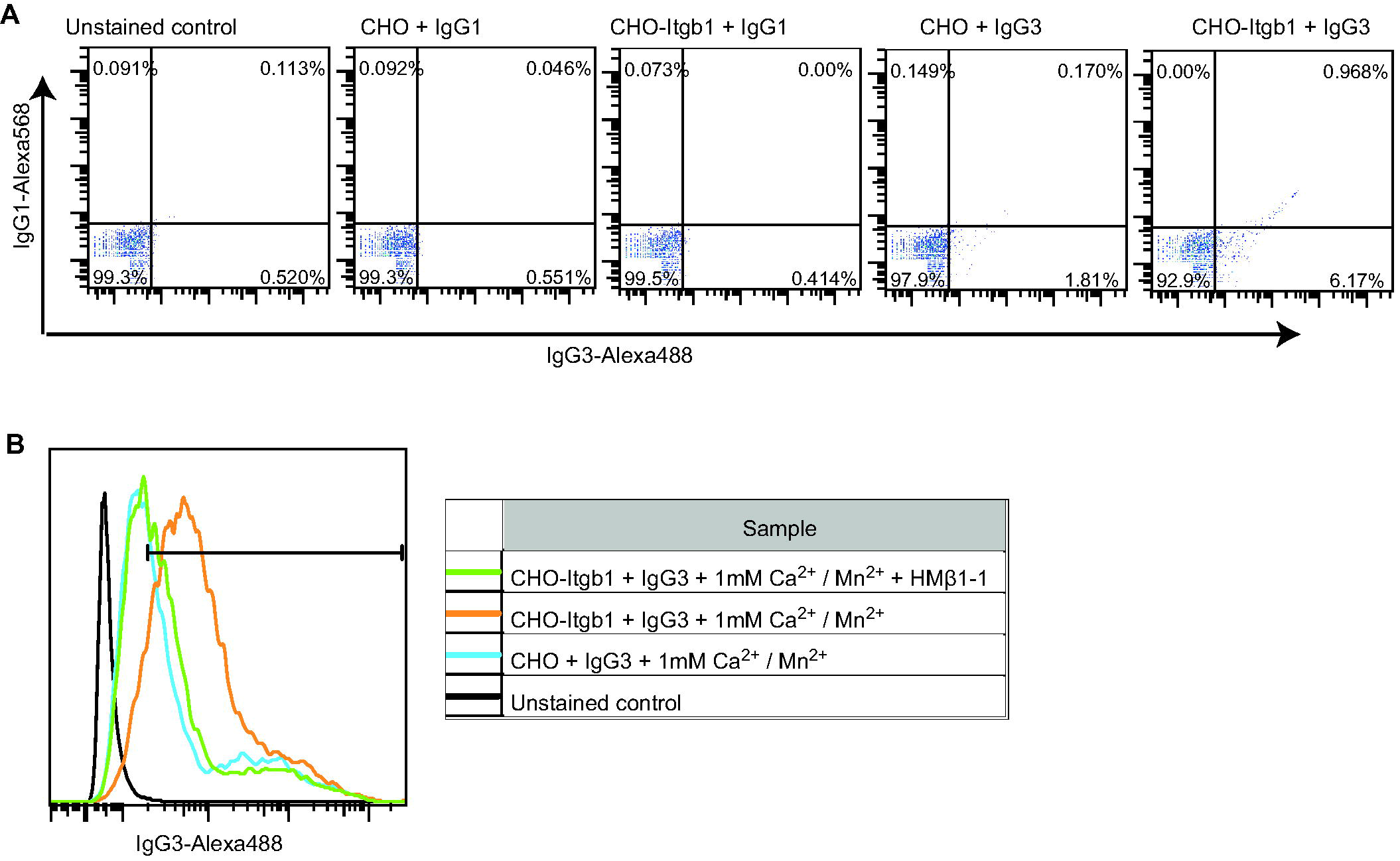
Expression of *Itgb1* in knockout cells confer binding to mIgG3. A) Fluorescently labeled 3E5-mIgG1 or 3E5-mIgG3 were incubated with untransfected CHO cells or CHO-Itgb1 cells, which express murine *Itgb1*. (B) Binding in the presence of 1 mM Ca^2+^ / Mn^2+^ and HMβ1-1.

### A new model for mIgG3-receptor interactions

Almost four decades ago Diamond and Yelton proposed the existence of another antibody receptor. Since that time, several research groups have delved into the issue of the mIgG3 receptor. Three different hypotheses surfaced from these studies:

#### Hypothesis 1 - Interactions between complement and mIgG3 explain function of mIgG3

In a model of anemia induced in mice by injection of high affinity anti-RBC antibodies, Fcγ knockout reduced mIgG2a-, mIgG2b-and mIgG1-induced but had no effect on mIgG3-induced anemia. However, the mIgG3 anemia was abrogated in C3-deficient animals, suggesting that it functions simply by activating complement (Azeredo da Silveira et al., 2002). A monoclonal mIgG3 to *Candida albicans* was also only protective when complement C3 was available (Han et al., 2001). Hjelm and colleagues also found that mIgG3 enhancement of the antibody response was normal in FcγRI-deficient mice, but that this enhancement was abolished in the absence of complement (Hjelm et al., 2005). Earlier, Stahl et al had also found that knocking out Fcγ receptors made no difference in mIgG3 enhancement and that C3 was important for IgG3-mediated antibody enhancement; however, they were not able to completely abrogate the enhancement and found lower but non-negligible enhancement with mIgG3 in animals depleted from C3 with cobra venom or in CR2 knockout mice (Diaz de Stahl et al., 2003).

#### Hypothesis 2 - FcγRI is the mIgG3 receptor

Experiments made with macrophages from transgenic mice showed that FcγRI knockout abolished mIgG3-mediated phagocytosis of erythrocytes.(Gavin et al., 1998). This hypothesis can be further supported by earlier studies with murine monoclonal antibodies to human CD3, which showed that human monocytes bound mIgG3 via the high affinity FcγRI (Ceuppens and Van Vaeck, 1989; Parren et al., 1992).

#### Hypothesis 3 - There is an unknown dedicated receptor for mIgG3

In a murine model of malaria, an mIgG3 monoclonal antibody remained protective in animals lacking the common Fcγ chain (Rotman et al., 1998) and the FcγRI alpha chain (Vukovic et al., 2000). Our group has shown evidence of functional differences between a monoclonal mIgG3 and its isotype switches in a murine model of acute lethal toxicity (Lendvai et al., 2000). Moreover, we have observed mIgG3-mediated phagocytosis in macrophages derived from animals lacking several genes encoding Fcγ and complement receptor subunits (Saylor et al., 2010).

These three hypotheses appear at first glance to be mutually exclusive: the experiments leading to conclusion (1) contradict (2) and (3); in some reports mIgG3 mediated function even when complement was absent or complement receptors blocked, contradicting (1); in others, blocking or knocking out FcγRs had no effect in mIgG3 functions, contradicting (2). However, our results suggest a model that can reconcile all three hypotheses (Figure 6). We propose that integrins play a part in mIgG3 binding and phagocytosis, as do FcγRs and complement. In some of our experiments, (e.g. Figure 3A), Itgb1 blockage drastically reduced but did not abolish mIgG3 function whereas FcγRII/III blockage had no effect; the same antibody batch was used for the flow cytometry binding assay shown in Figure 2G, which shows excess soluble mIgG2a (a competitive FcγRI block) partially decreasing mIgG3 binding. Other antibody batches, such as the recombinant 2H1 antibodies used in the phagocytosis assay shown in Figure 3D, do show a similar pattern of mIgG3-mediated phagocytosis reduction upon Itgb1 blockage; the extent of the reduction, however, is not as large as what we observed with 3E5-mIgG3. Moreover, FcγRII/III blockage did affect 2Hl-mIgG3-mediated phagocytosis, whereas it did not affect 3E5-mIgG3. Adding up observations of different mIgG3 batches mediating phagocytosis by integrin and FcγR to different extents, the partial competition with monomeric mIgG2a and the fact that in none of the experiments integrin blockage completely abolished mIgG3 function or binding suggests that monoclonal mIgG3 preparations are actually a mixture of isoforms binding to FcγRs and to Itgb1. The existence of a mixture of isoforms that varies in each batch could explain the seemingly disparate data on whether FcγRI or an unknown protein is the mIgG3 cell surface receptor. Such heterogeneity is very common and has been extensively studied in industrial monoclonal antibody production settings, in which variations in glycosylation have significant impact on biologic drug pharmacokinetics and pharmacodynamics (Liu, 2015).

**Figure 6.**
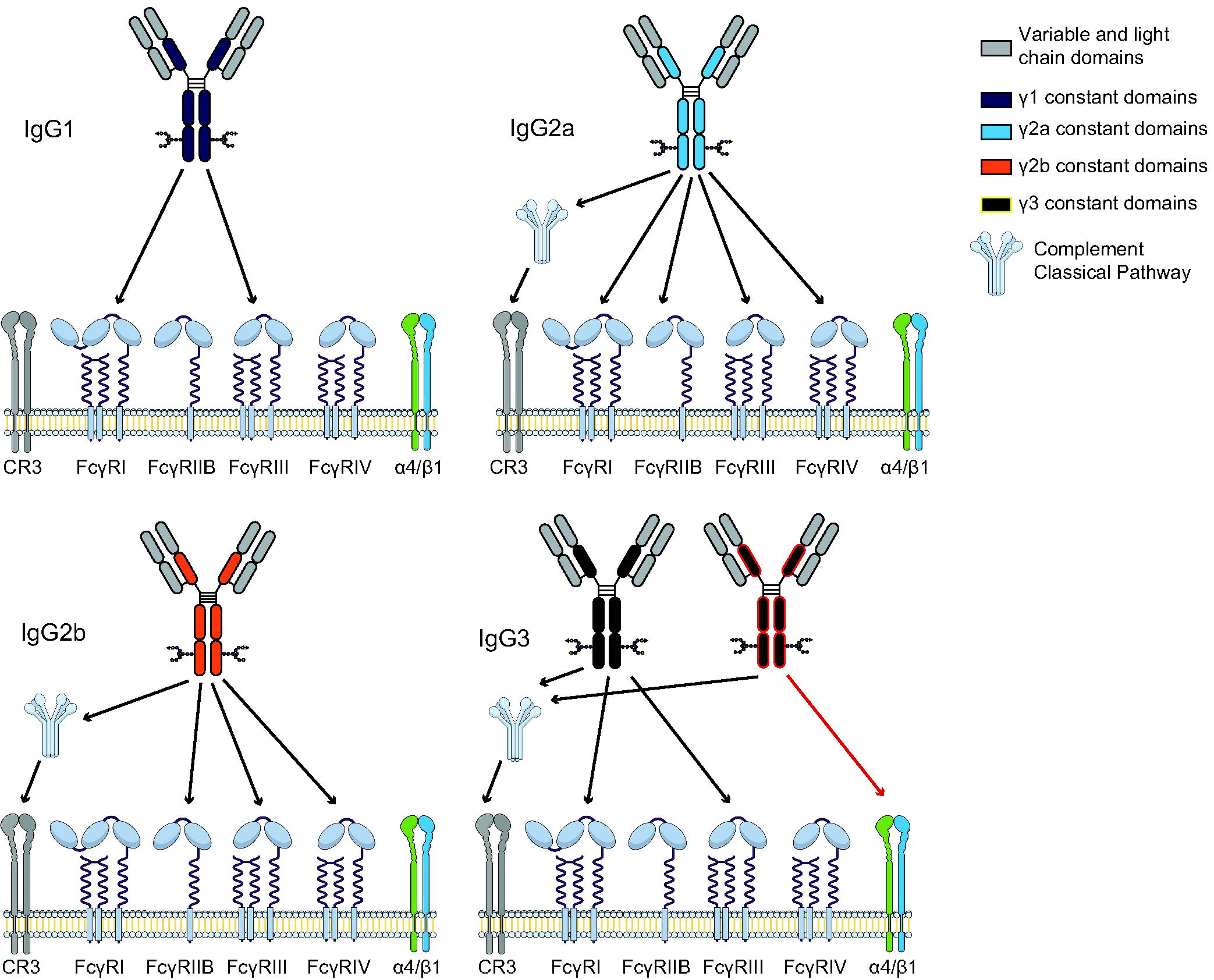
Model for mIgG interaction with cell surface receptors. Model summarizing the cell surface receptors that mediate mIgG functions in macrophages. Black arrows indicate known functions, including the direct interaction between mIgG antibodies and Fcγ receptors (Bruhns, 2012; Nimmerjahn et al., 2015) and the indirect interaction with complement receptor, mediated by classical pathway activation. Red arrows indicate the Itga4/Itgb1 role in mIgG3 function described in this manuscript.

## Discussion

Antibody isotype switching in response to antigens of different compositions or origins is conserved in many animals, probably because different antibody isotypes have different effector functions and thus allow a host to tailor the immune response to each particular threat. Antibody engineers also change the isotype of the antibodies they work with to obtain biotechnological or pharmaceutical tools with different effector functions. In both cases, most of the functional differences between isotypes can be ascribed to their differential interactions with cell surface receptors.

Integrins are transmembrane receptors present as α + β heterodimers that regulate cell-cell and cell-matrix interactions (Cabodi et al., 2010). Several integrins also have an intrinsic ability to mediate phagocytosis as well (Dupuy and Caron, 2008). Probably the most studied ones are the β2 integrin phagocytic complement receptors CR3 (αM + β2) and CR4 (αX + β2). In the *C. neoformans* system, Itgb2 was shown to mediate complement-independent phagocytosis as a result of direct interactions with the capsular polysaccharide component of antibody-coated cryptococcal cells (Taborda and Casadevall, 2002). β1 integrins can also mediate particle internalization: (1) macrophages lacking Itgb1 expression manifest reduced phagocytic capacity for some types of bacteria (Wang et al., 2008); (2) microglia are able to phagocytose fibrillar beta-amyloid in vitro in a mechanism that does not utilize FcγRs or CRs and is blocked by an antagonist to Itgb1 (Koenigsknecht and Landreth, 2004); and (3) Itgb1 pairs with α2 integrin to form a receptor that binds complement component C1q (Zutter and Edelson, 2007). The interaction between IgG molecules and non-FcγR cell surface proteins has also been reported before (Anthony et al., 2011).

Additional evidence for the conclusion that Itgb1 is involved in mIgG3 function comes from experiments using divalent cations. Integrins have many binding sites for these ions, each with distinct functional roles (Zhang and Chen, 2012). For most integrins, Mn^2+^ binding to one of these motifs (the metal ion-dependent adhesion site - MIDAS) changes the integrin conformation into an active open state. Consistent with this, Mn^2+^ increased binding of IgG3 to surface of cells. Mg^2+^, which also binds to MIDAS to activate integrins, resulted in more mIgG3 bound to J774 cells; however, the activation mediated by Mg^2+^seemed to be different from the Mn^2+^ activation, as addition of Mg^2+^ competed with Mn^2+^ and completely abolished increased mIgG3 binding. In addition to the Mn^2+^/Mg^2+^ MIDAS, integrins have different types of Ca^2+^ binding sites (Leitinger et al., 2000). One of them, with mM affinity, flanks MIDAS and, when occupied, inhibits Mg^2+^/Mn^2+^ binding and thus integrin function. However, Ca^2+^ actually increased mIgG3 surface binding in our assays. This seemingly surprising result is actually very informative, as a detailed study on the effect of calcium in β1 integrins showed it decreased the activation of the heterodimers containing α2, α3, α5 and α6 but increased the activation of the α4/β1 integrin (Bazzoni et al., 1998), an indication that Itga4 is the Itgb1 partner in the mIgG3 receptor.

Given that the problem of the mIgG3 receptor identity has vexed investigators and lingered unsolved for decades, we arrive at our conclusions humbly. The most straightforward interpretation of our findings is that Itga4/Itgb1 is at least part of an mIgG3 receptor complex. We acknowledge that we lack direct evidence for an immunoglobulin-integrin protein-protein interaction, which in combination with the functional data from our study would provide unequivocal evidence for an mIgG3 integrin receptor. Nevertheless, the available data provide strong evidence associating integrin function with IgG3 phagocytosis, which leads to a tantalizing suggestion regarding the evolution of the humoral response. Antibodies appeared around half a billion years ago, with cartilaginous fishes. The Fcγ receptors, however, are much more recent: they first appeared with mammals. The most accepted interpretation is that through most of its existence, antibodies functioned solely by activating complement and neutralizing antigens. However, as integrins are far more ancient and are present in all metazoans, they could actually have functioned as early IgG receptors for non-mammalian vertebrates. In this regard, it is interesting to note that mIgG3 is the murine IgG isotype whose sequence differs the most from the others and which seems to have appeared first in the evolutionary history of the mouse constant gamma locus (Hayashida et al., 1984).

## Acknowledgements

We would like to acknowledge Dr. Jinghang Zhang from the Albert Einstein College of Medicine flow cytometry core facility for technical assistance. AC is supported by National Institutes of Health Grants 5R01A1033774, 5R37AI033142, and 5T32A107506, and CTSA Grants 1 ULI TR001073-01, 1 TLI 1 TR001072-01, and 1 KL2 TR001071 from the National Center for Advancing Translational Sciences. Facilities were supported by the Center for AIDS research at Albert Einstein College of Medicine; CC was partially supported by PhD grant SFRH / BD / 33471 / 2008 by Fundação Ciência e Tecnologia; CS was supported by NIH grant T32 AI07506; AN was partially supported by a Reuni scholarship (MEC-Brasil) and is currently supported by grants from the Brazilian funding agencies CNPq, FAP-DF and Capes.

## Materials and Methods

### Mice

Wild-type C57Bl/6 mice (Jackson Laboratories, Bar Harbor, ME), from colonies maintained at the Animal Facility of Albert Einstein College of Medicine, were used to obtain the peritoneal macrophages. To obtain Itgb1-deficient macrophages (*Itgb1-/-*), a conditional knockout model had to be used because the Itgb1 knockout is lethal. Mice with loxP sites flanking exon 3 of the *Itgb1* gene (Raghavan et al., 2000) were crossed with mice expressing the Cre recombinase under control of the lysozyme promoter (Clausen et al., 1999). In all experiments, mice were treated in accordance with institutional guidelines, and the animal protocols were accepted by the institutional review committee of the Albert Einstein College of Medicine, where the experiments were made.

### Microbe strains

*Cryptococcus neoformans* strains 24067 (serotype D) and H99 (serotype A) were used. The yeast cells were grown overnight in Sabouraud dextrose broth (Difco) at 30°C with agitation.

### Cell lines

For experiments the following cell lines were used: J774.16 (J774) - murine macrophage-like cells; L929 - murine fibroblast, producer of macrophage colony stimulating factor (M-CSF); CHO-K1 - Chinese hamster ovary cells; A549 - human alveolar epithelial cells; GD25 - murine differentiated stem cells with Itgb1 knockout (GD25−/−) and reconstitution (GD25B1A) (Fassler et al., 1995); NSO - murine myeloma, used to produce recombinant antibodies; 293F - suspension-adapted human embryonic kidney cells. J774, L929, CHO-K1, A549 and GD25 cells were grown at 37°C and 5-10% CO_2_ and maintained in Dulbecco’s Modified Eagle’s Media (DME) supplemented with 10% fetal bovine serum (from North American or South American origin) that was heat-inactivated by treatment at 56°C for 30 min, 10% Gibco NCTC-109 Media (Invitrogen), 1% non-essential amino acids (Mediatech or Gibco) and 1% Penicillin Streptomycin Solution (Mediatech or Gibco). NSO cells were maintained in the same supplemented DME medium and were grown in CD Hybridoma chemically defined medium (Invitrogen) in a 37°C incubator with 125 rpm shaking and 5% C0_2_. 293F cells were also grown in suspension of Freestyle 293F medium (Invitrogen) also at 37°C, 5% C0_2_ and 125 rpm shaking. Adherent cells were grown on tissue culture treated plates (BD Falcon) and removed by treatment with 3-5 mL Cellstripper (Mediatech), a non-enzymatic solution for removing adherent cells, or 5 mL trypsin (Gibco) then pelleted by centrifugation at 300 x g for 10 min, and finally re-suspended at the appropriate concentration in DME.

#### Primary cell cultures

Bone marrow-derived macrophages (BMMs) were differentiated from the marrow of femoral and tibial bones of donor mice. Briefly, cells were obtained by flushing the marrow and then growing in differentiation media for 6 d (DME media with 20% conditioned medium from a confluent culture of L929 fibroblasts as a source of M-CSF, 10% FCS, 10 mM Hepes, 2.0 mM L-glutamine, 0.05 mM 2-mercaptoethanol, 1% nonessential amino acids, 1% penicillin and 100 μg/ml streptomycin). Nonadherent cells were removed and adherent macrophages recovered from plates using Cellstripper. These BMMs were then maintained in DME media with 10% L929 conditioned medium, 10% FCS, 10 mM Hepes, 2.0 mM L-glutamine, 0.05 mM 2-mercaptoethanol, 1% nonessential amino acids, 1% penicillin and 100 μg/ml streptomycin, and used within 3 days.

### Method details

#### Generation of genetically modified cell lines

We initially amplified murine *Itgb1* and *Itga4* from J774 total RNA by RT-PCR. To generate Itgb1+ CHO cells, the *Itgb1* insert was cloned on a pEGFP-C1 vector (Clontech, Mountain View, CA) that was transfected into CHO cells using Lipofectamine 2000 (Invitrogen, Carlsbad, CA). Stable transfectants were selected with 1 mg/mL G418 and a population with high murine *Itgb1* expression was selected by two consecutive rounds of flow cytometry sorting using PE-labeled HM β 1-1. To generate GD25 cells expressing *Itgb1, Itga4* or both integrins, the *Itgb1* insert was cloned in the pIRES2-EGFP-blas vector (in which the NeoR resistance marker was substituted by the BlasR marker by Infusion cloning) and the *Itga4* insert was cloned into the pIRES2-DsRed2 vector. Pairwise combinations of the *Itgb1*- and *Itga4*-expressing plasmids as well as the empty vectors were co-transfected into GD25−/− cells using Lipofectamine 3000 (Invitrogen). All the sequences in these vectors were confirmed by Sanger sequencing before transfection.

#### Hybridoma antibodies

The *C. neoformans*-specific monoclonal antibodies, 3E5 IgG1 and IgG3 have been described previously (Yuan et al., 1995). Briefly, mAb 3E5 was originally isolated as an IgG3 mAb following immunization of mice with GXM conjugated to tetanus toxoid (Mukherjee et al., 1993), and the other isotypes were later generated via *in vitro* isotype switching (Yuan et al., 1995). Ascites was generated by injecting hybridoma cells into the peritoneal cavity of pristane-primed BALB/c mice (National Cancer Institute, Frederick, MD) and harvesting the fluid. Abs were purified from ascites using a protein G column following the manufacturer’s instructions (Pierce, Rockford, IL), then dialyzed in PBS and quantified by ELISA with an isotype-matched standard to determine concentration. For quantification, ELISA plates were coated with serial dilutions of purified myeloma IgG1/k (MOPC 21) and IgG3/k (FLOPC 21) standards (Cappel) and dilutions of the purified antibodies. After blocking with bovine serum albumin (BSA), bound antibodies were detected with alkaline phosphatase-conjugated isotype-specific goat antimouse polyclonal serum (Southern Biotech).

#### Recombinant antibodies

##### Synthetic genes and cloning

To produce 2H1 and 4-4-20 mIgG1/mIgG3 recombinant antibodies, the heavy (VH) and light chain (VL) variable regions of 2H1 (Young et al., 1997) and 4-4-20 (Whitlow et al., 1995) antibodies were codon-optimized and synthetized by GenScript and then cloned into the commercially available pFUSE vectors, the VHs into the pFUSEss-CHIg-mG1 and pFUSEss-CHIg-mG3 vectors and the VLs into the pFUSEs-CLIg-mk (InVivoGen). All constructions were confirmed by Sanger sequencing before transfection. NSO cells were seeded at 8 × 10^5^ cells/mL in 24-well plates 24 hours before transfection and then co-transfected with the pFUSEss-CHIg-mGl or pFUSEss-CHIg-mG3 and the pFUSEs-CLIg-mk vectors containing the VH and VL sequences for 2H1 or 4-4-20 antibody expression. Transfections were made using Lipofectamine2000 (Invitrogen), following manufacturer instructions. 72 hours post-transfection cells were set to selection by addition of Zeocin (Invitrogen) at 1 mg/mL and Blasticidin (Gibco) at 5 μg/ml. After three weeks of selection stable antibody producing cells were obtained. These cells were adapted to serum free medium, CD Hybridoma AGT (TermoFisher), supplemented with 1x cholesterol (Gibco) and 8 mM L-Glutamine (Sigma Aldrich). The culture volume was then escalated to 1 L to produce the first lots of 2H1 and 4-4-20 mIgG1 /mIgG3 recombinant antibodies. Subsequent lots of the recombinant antibodies were produced by thawing and expansion of frozen antibody producing cells adapted to serum free medium, obtained as described above. Recombinant antibodies were purified by either, anionic exchange chromatography using a HiScreen DEAE FF column (GEHealthcare) in a non-binding mode, followed by a two-step ammonium sulphate precipitation at 64% saturation, size exclusion chromatography with PBS buffer using a HiLoad 16/600 Superdex 75 pg column (GEHealthcare), or by affinity chromatography using Protein G GraviTrap columns (GEHealthcare), following instructions on manual, and then concentrated by ultrafiltration. Quantification was made by direct ELISA as described above for hybridoma antibodies or by an antigen-specific ELISA for preparations with low concentration.

For the antigen-specific ELISA, plates were coated with 10 μg/mL of GXM purified from *C. neoformans* H99 cultures. After blocking with bovine serum albumin (BSA), serial dilutions of isotype specific antibodies IgG1 (18B7) or IgG3 (3E5) and purified recombinant antibodies were added to wells and detected with alkaline phosphatase-conjugated isotype-specific goat anti-mouse polyclonal serum (Southern Biotech). 2H1 antibodies were validated by indirect immunofluorescence with *C. neoformans* cells (Figure S4).

#### *C. neofomans* phagocytosis assay

Phagocytosis assays were performed in 96 well tissue-culture treated plates (BD Falcon) containing either primary peritoneal cells isolated one day prior to the experiment, BMM cultured to maturity, or J774 cells plated at least 2 h or up to one day before the experiment. In some experiments, various receptors were blocked using receptor-specific Abs prior to phagocytosis. For CR block, a cocktail containing Abs to CR3 and CR4 (CD18, CDllb, and CDllc; BD Pharmingen) are known to be effective at blocking CR function (Taborda and Casadevall, 2002). For FcγRII and FcγRI 11 block, the 2.4G2 Ab (BD Pharmingen) was used. For FcγRI block, monomeric IgG2a was used as a competitive inhibitor. For Itgb1 block, the unlabeled mAb HMβ1-1 was used, which has been shown to be effective at binding and inhibiting adhesion of tumor cell lines to extracellular matrix via Itgb1 (Noto et al., 1995). Blocking Abs were added at concentrations ranging from 10 to 50 μg/mL 30 minutes before addition of the fungi and opsonizing antibody. Then, the *C. neoformans* suspension with opsonizing Ab (IgG1, IgG3 or 18B7, as indicated) was added, with blocking Abs at 10 μg/mL, opsonizing Abs at 10 μg/mL, and the macrophage to *C. neoformans* ratio ranging from 1:1 to 1:2. Phagocytosis was allowed to proceed for 2 h, at 37°C in 5-10% CO_2_. Cells were then washed, fixed with methanol at −20°C for 30 minutes and finally stained with Giemsa. Cells were then analyzed under an inverted microscope, counting three fields/well, with at least 100 cells/field. Macrophages with internalized *C. neoformans* were readily distinguishable from cells that had taken up no fungi, or from those in which *C. neoformans* was simply attached to the outside, due to the visible vacuole containing engulfed cell. Percent phagocytosis is calculated as the number of macrophages containing one or more internalized *C. neoformans* divided by the total number of macrophages visible in one field. At least one hundred macrophages were counted in each condition.

Phagocytosis assays to determine alpha integrin partner were performed using an automated microscopy platform, as described previously (Coelho et al., 2012). Briefly, macrophages were seeded in a glass-bottom 96-well plate (MGB096-1-2-HG-L; Matrical Biosciences, Spokane, WA) at a density of 2.4×l0^4^ cells/well and phagocytosis of *C. neoformans* proceeded as described above. After a 2 h period to allow phagocytosis, cells were fixed with methanol and stained consecutively with wheat germ agglutinin conjugated to Alexa Fluor 633 (Invitrogen) for detection of macrophage cytoplasmic membrane, Uvitex 2B (Polysciences, Inc., Warrington, PA) at a 0.1 g/mL to stain *C. neoformans* cells and finally propidium iodide (Sigma) at 5 μg/mL. Images were acquired and analyzed in iCys Compucyte (CompuCyte Corp., Westwood, MA) with a 40x objective with at least 15 images per field averaging around 700 cells per well in triplicate wells.

#### Pooled shRNA library transduction and sorting

A pool of lentiviral vectors containing approximately 80,000 different shRNA sequences targeting about 15,000 different mouse genes was obtained from Sigma-Aldrich, the distributor for the product developed by the RNAi Consortium (Luo et al., 2008; Root et al., 2006). J774 cells were maintained as indicated above and were plated in 10 cm tissue cultured treated plates (BD Falcon) one day prior to transduction to generate 10 plates the next day, each at a density of 2 × 10^6^ J774 cells per plate. Hexadimethrine bromide (Sigma) was added at 8 μg/mL to facilitate transduction, and 2 × 10^5^ viruses were added to each plate of J774 cells to maintain a multiplicity of infection of 0.1. Cells were incubated with virus 24 h, the media was then changed and the cells were incubated an additional day. At this point, 5 μg/mL of Puromycin (Sigma) was added and cells were grown under selection for 2 d. After selection, dead cells were removed and cells were replated under continued puromycin selection, and allowed to recover overnight. Next day the 4.6 × 10^7^ total cells were stained with two labeled Abs: mIgG3-DTAF and mIgG1-APC. The mIgG3 was the 3E5 mAb featured in other experiments, generated as indicated in the previous section, and labeled with dichlorotriazinylaminofluorescein (DTAF, Sigma) according to the protocol for labeling amine-reactive probes following provided instructions (Invitrogen). Mouse IgG1 isotype control Ab conjugated to APC was purchased (Southern Biotech). For the sorting of transduced cells: staining was carried out in Cellstripper for 1 h at 4°C. Afterwards, the cells were collected by centrifugation, washed twice, and then filtered in a 40 μm cell strainer and analyzed on the MoFlo XDP Sorter (Albert Einstein College of Medicine Flow Cytometry Core Facility). Sorting gates were generated for cells with high IgG1 signal and low IgG3 signal, and this first sort was run in “yield” mode to get the highest return. We recovered 1.5 × 10^6^ cells that were then grown under continued puromycin selection for one week, after which we performed a second sort under similar conditions as above except in “purity” mode to increase the selectivity of the method and recover only those cells in the sorting gate population. We recovered approximately 50,000 cells in the post-sort population and maintained them in culture under puromycin selection. A portion of J774 cells not subjected to sorting was also maintained as the pre-sort population. These cell populations were maintained under puromycin selection and used for DNA extraction and sequencing. Following bioinformatics analysis, 17 specific shRNA lentiviral vectors were purchased, corresponding to our candidate selection criteria. These individual shRNAs were transduced into J774 cells following a similar protocol, maintained as separate transduced cell lines, and analyzed as indicated.

#### Flow cytometry and sorting

To analyze the binding of either IgG1 or IgG3 mAbs, or to analyze expression of Itgb1, transduced J774 cells were maintained as indicated above and prepared for flow cytometry. Approximately 1 × 10^6^ cells were harvested and suspended in 1 ml Cellstripper solution to prevent attachment, and 10 μl IgG1-Alexa568 and IgG3-Alexa488 were added. These Abs were made by conjugating the 3E5 IgG1 and IgG3 mAbs with their respective Alexa fluorophore following the kit instructions (Molecular Probes, Eugene, OR). For staining Integrin beta 1 (Itgb1), phycoerithrin-labeled HMβ1-1 was used at 1:100 dilution. After staining for 1 h at 4°C, cells were centrifuged and washed twice, then analyzed on BD LSR II or BD FACS Aria at the Albert Einstein College of Medicine Flow Cytometry Core Facility. Flow cytometry data were analyzed with Flow-Jo (Treestar, Inc.) and presented as the median fluorescence intensity, histograms or density plots. For the case of Itgb1 sorting, J774 cells transduced with the 5 individual shRNAs targeting Itgb1 were analyzed for surface expression, and the group of cells with lowest Itgb1 staining were sorted and maintained.

#### DNA extraction and sequencing

Genomic DNA was extracted from the pre-and post-sorted J774 cells populations using a commercial kit (QIAGEN, Valencia, CA). Primers flanking the shRNA region were used to amplify the shRNA sequences from 1.5 μg of DNA. The amplified library was then analyzed with a Solexa sequencer (Illumina, Inc., San Diego, CA). Partners Healthcare (Cambridge, MA) performed PCR amplification and high-throughput sequencing on pre-and post-sort DNA samples. Seqwise LLC (Boston, MA) performed bioinformatics on the sequencing data to generate actual counts.

### Quantification and statistical analysis

Bioinformatics data from the shRNA high-throughput sequencing was analyzed with the Audic and Claverie method for pairwise testing (Audic and Claverie, 1997; Romualdi et al., 2003). Phagocytosis assays were analyzed on Graphpad Prism 7.0; pairwise comparisons were made with the Fisher’s exact test to compare the proportions of macrophages which internalized *C. neoformans* cells. Confidence intervals for the proportions were calculated using the Wilson/Brown method.

